# miR-17∼92 Suppresses Proliferation and Invasion of Cervical Cancer Cells by Inhibiting Cell Cycle Regulator Cdt2

**DOI:** 10.1101/2022.12.30.522306

**Authors:** Garima Singh, Sonika Kumari Sharma, Aastha, Samarendra Kumar Singh

## Abstract

Cervical cancer (CC) is 4^th^ largest killer of women worldwide when diagnosed in late stages the treatment options are almost negligible. 99% of CC is caused by high risk human papilloma viruses (HR-HPV). Upon integration into human genome the encoded viral proteins mis-regulates various onco-suppressor and checkpoint factors including cell cycle regulators. One such factor is cell cycle S phase licensing factor Cdt2, which has been reported to be highly upregulated in various cancers including CC. HPV proteins also suppress several tumor suppressor miRNAs concluding miR-17-92 cluster. In this study we report that miR-17-92 directly recruits to 3’UTR of Cdt2 and downregulates this oncogene which suppresses the proliferation, migration, invasion capabilities of the CC cell lines while normal cells are fine. Suppression of Cdt2 by miR17-92 blocks the cancerous cells in S phase and induces apoptosis eventually leading to their death. Hence, our work for the first time mechanistically shows how miR17-92 could work as tumor suppressor opening up the potential of miR17-92 to be used in developing therapy for cervical cancer treatment.

## Introduction

Cervical cancer (CC) is one of the most lethal diseases caused by Human Papilloma Virus (HPV) with a survival rate of approximately 15% when diagnosed in late stage^1,2^. It is the fourth most leading cause of death among women world-wide with approximately 604,000 new cases and 342,000 deaths reported annually which is estimated to increase by 50% till 2030^3^. HPV encoded oncoproteins E5, E6 and E7 interacts with several cellular factors and drives them to mis-regulation causing carcinogenic transformation of the healthy cells. The transformed cells have suppressed apoptotic checkpoint system including p53, pRb and several others resulting in hyper-proliferation of the cancerous cell^4,5^. The E6 protein of HPV alone works at multiple genes to inhibit apoptotic signaling pathway and activates telomerase reverse transcriptase to promote immortalization of cells. Apart from this, E6 also leads to ubiquitin mediated degradation of guardian of genome i.e., p53 protein via E6AP ubiquitin ligase system^6^. E6 stabilizes a major cell cycle regulator protein CDC-10 dependent transcript-2 (Cdt2)/DTL by recruiting a de-ubiquitinase USP46, which was discovered to be essential for proliferation and survival of the cancer cells^7^. Cdt2 protein level has been reported to be highly amplified in various cancers including cervical cancer^8^.

Cdt2/DTL/DECAF2 is an essential component which was first discovered in fission yeast^9^. It is a substrate receptor DECAF (Ddb1-and Cul-4 associated factor) for CRL4 Cullin RING E3 ligase system, to form the complex CRL4^Cdt2^ which functions as a master regulator of cell cycle progression and genome stability^10^. CRL4^Cdt2^ E3 ubiquitin ligase ensures the timely degradation of various cell cycle factors like p21, Set8 and Cdt1 etc. which are involved in DNA replication initiation, apoptotic checkpoint regulation and chromatin modification to prevent re-replication in S phase and is also required for the early G2/M checkpoint^11,12^ system. Like all the other cell cycle regulators, the level of Cdt2 protein is also maintained by several ubiquitination systems like CRL1^FBXO11^, CRL4^DDB2^, APC/C-Cdh1 etc. which leads to its elevated expression levels during G1 to S phase transition and decreased levels during mitosis^8^.

Even after the availability of vaccines against HPV (16 and 18), due to the lack of awareness and proper planning in low and middle-income countries (accounting for approximately 90% of cases), there has not been any impactful decline either in cervical cancer cases or in mortality rates^13,14^. The standard treatment available to cure cervical cancer is radiotherapy in combination with chemo and brachytherapy to which most of the patients do not respond and have high rate of relapse and lower survival rates. Therefore, we need a therapy which could be effective enough to majority of population. In this direction, miRNAs have emerged as target specific and effective therapeutic agents which could prove to be effective in regulating cancer progression.

MiRNAs are small non-coding RNAs (17-25 nucleotides in length) which play diverse roles in controlling gene expression by degradation/translation repression of target mRNAs. MiRNA works by targeting 3’UTR (Untranslated Regions) sequences of mRNA which are complementary to their seed sequence of 6-8 base pairs^15^. Each miRNA can regulate several (hundreds) mRNAs and are involved in regulating major processes like cell cycle, proliferation, apoptosis, development etc. and approximately 60% of human genes are estimated to be regulated by miRNAs^16^. These miRNAs can either be encoded as a single miRNA or as a cluster of multiple miRNAs. In vertebrate genome, about 30% of miRNAs occur in the polycistronic cluster^17,18^. miR-17∼92 also known as oncomiR-1, is one of the polycistronic cluster, located in MIR17HG (miR17-92 cluster host gene; non-protein coding) intron on chromosome 13 (13q31.3), encoding six mature miRNAs: miR-17, miR-18a, miR-19a, miR-20a, miR-19b-1, and miR-92a-1^19–21^ (Figure.1). Expression of miRNA-17∼92 is essential for development, regulation of cell cycle phenomenon, proliferation, and other several vital cellular functions. Dysregulation of the same has been found in various cancers and diseases and is linked to their progression^22^. This cluster of miRNAs is regulated by the direct interaction of two transcription factors, cellular myelocytomatosis oncogene (c-Myc) and n-Myc at the promoter region of miR-17∼92 to initiate its transcription^23–25^. The interaction of miR-17∼92 and these transcription factors works in autoregulatory feedback loop, where two members of miR-17∼92 cluster; miR-17, and miR-20a regulate the translation of E2F1-3 which further regulate the expression of c-Myc to maintain the cellular balance between apoptosis and proliferation of the cell^23,26,27^ (Figure.1). The studies till date have shown miR-19a and miR-19b as the key oncogenes whereas miR17 and miR18a act majorly as tumor suppressors in the cluster^20,21^ where miRNA17 acts by targeting CCND2 (cyclin D2)^28^ and miR-18a affects miR17-92 biogenesis by regulating expression of Dicer enzyme (a member of miRNA biogenesis pathway). Apart from this, miRNA-18a directly targets the tumor protein TP53-regulating inhibitor of apoptosis gene 1 (TRIAP1) and inositol phosphate multi-kinase (IPMK)^19^. Various studies have established both the oncogenic and tumor suppressor role of the miRNA17-92 cluster and the role of the individual miRNAs in this cluster remains unclear^19–21^. We became interested in miR-17∼92 while looking for the effect of miR-34a on expression of Cdt2 protein^15^. In early screening, we discovered that miR-17∼92 cluster suppresses Cdt2 expression level in cervical cancer cell lines. We further explored the effect of miR-17∼92 on proliferation, invasion and metastasis behavior of cervical cancer cells and discovered that it suppresses both proliferation and migration of these cells. This is the first ever to report that miR-17∼92 interacts with Cdt2 (at 3’UTR) and inhibit both at transcript and protein level. Our finding could pave the way to explore the possibility of using miR-17∼92 as miRNA therapy to manage cervical cancer.

**Figure 1:**
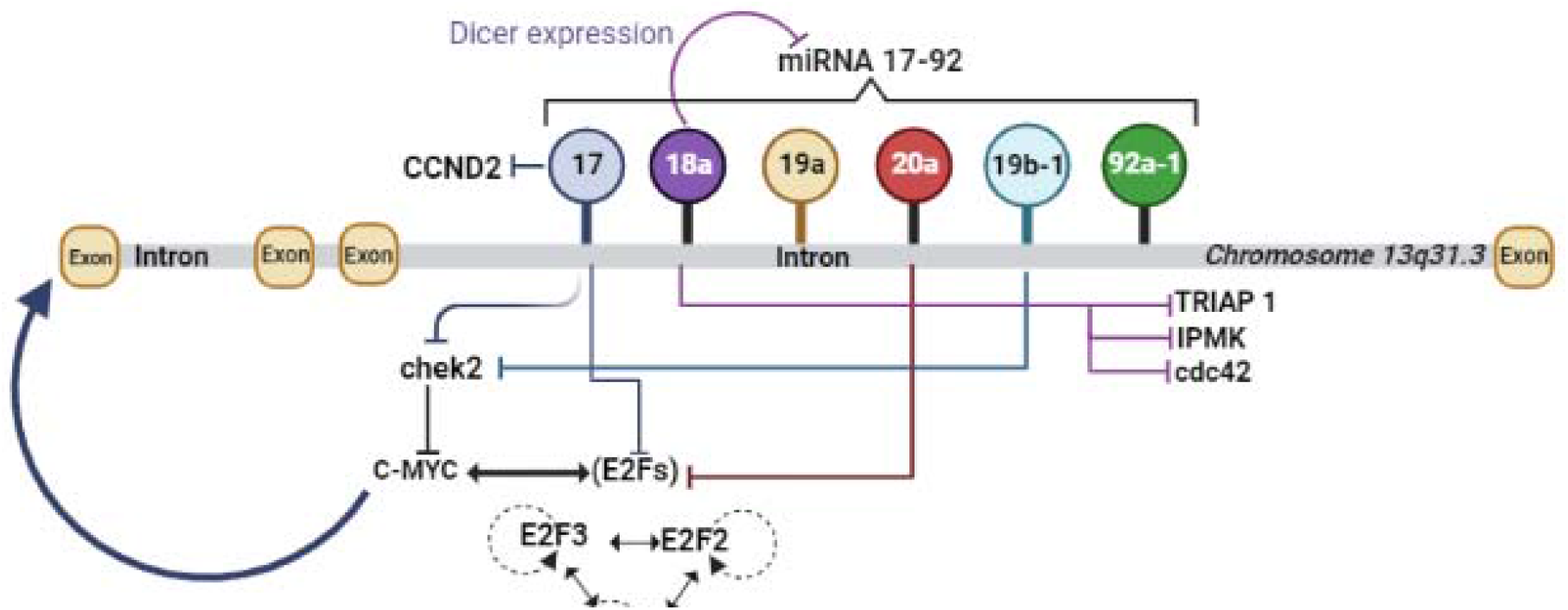
Coordinated transcriptional activation and the autoregulatory feedback loop of miR-17∼92 (Created in BioRender.com).

## Materials and Methods

### Cell Lines and Media

SiHa and HeLa cell lines (HPV positive cervical cancer cell lines) were purchased from the American Type Culture Collection (ATCC, USA). C33A (HPV negative cervical cancer cell line) and HEK293T (Human kidney cell line) were procured from National Centre for Cell Science (NCCS, Pune, India). The cells were cultured in Dulbecco’s Eagle Modified medium (DMEM, Gibco, USA) supplemented with 10 % Fetal Bovine Serum (FBS, Gibco, USA) and 1% Pen-Strep (Penicillin-Streptomycin, Gibco, USA) unless mentioned otherwise and cultured at 37□temperature, 95% humidity and 5% CO□.

### Plasmids and miRNA

pcDNA 3.1 plasmid vector acquired from Addgene (Watertown, USA) was used as a control. miRNA-17∼92 cloned in pcDNA 3.1 plasmid was obtained from Joshua Mendell lab^23^. Flag-Cdt2 plasmid was received as a gift from Dutta’s Lab (University of Virginia, USA). Rest of the miRNA mimics were purchased from Dharmacon (USA).

### Cell Transfection

24 hours before transfection, 0.1 million cancerous (HeLa, SiHa, C33A) and non-cancerous cells (HEK293T) were seeded in 6 well plates. The cells were then transfected with the plasmid vectors (pcDNA 3.1, miR-17∼92 (4 μg)), Flag-Cdt2 (2 μg)) with turbofect™ (Thermo Fisher Scientific, Massachusetts, USA) according to the manufacturer’s protocol. The cells were then cultured as mentioned above.

### Cell Proliferation Assay

After transfection of cells with miR-17∼92 and respective control plasmids, they were harvested at consecutive days (1,2,3,4 days). The harvested cells were then counted in triplicates, using trypan blue exclusion method (0.4% trypan blue solution) in the haemocytometer, as well as in cell counter (TC-20, Biorad). The growth curve was plotted and the significance of treatment was calculated using unpaired student’s t-test.

### Cycloheximide Half-Life Estimation Assay

SiHa/ HeLa cells were transfected with control (pcDNA 3.1) and miRNA-17∼92. After 24 hours of transfection freshly prepared cycloheximide (CHX, 25 μg/ml) was added to the culture. The cells were harvested at different time intervals (0,1,2,3 hours) followed by cell lysate preparation and western blotting.

### Western Blot Analysis

The cells were harvested and lysed using radioimmunoprecipitation assay (RIPA) buffer supplemented with PMSF (1mM) and protease inhibitor cocktail (Cell Signaling Technology, Massachusetts, USA). The protein concentration was determined using Bradford protein assay (Sigma, Missouri, USA). The total protein was then resolved on SDS-PAGE (8-12%) and transferred onto PVDF membrane (Millipore, Massachusetts, USA). After the transfer, using protein ladder as reference membranes were cut according to the protein of interest (as shown in the results). The further western blot protocol was followed as standard procedure^15^ and **e**nhanced chemiluminescence (ECL) substrate (Bio-Rad, California, USA) was used to develop the blot. Documentation of the blots was done using chemidoc system (Azure Biosystems 600).

### Cell Invasion Assay

After 48 hours of transfection, the cells (.1 million) were washed with PBS and resuspended in serum-free DMEM. Cells were then added onto the precoated upper chamber (with 2 mg/ml ECM gel; Sigma-Aldrich, Missouri, USA) of transwell plate (Corning, New York, USA), and in lower chamber DMEM supplemented with 10% FBS was added. Cells were allowed to invade for 48 hours in CO□incubator. Cells that remained in the upper chamber were removed whereas the cells which invaded through the membrane were first fixed with 5% glutaraldehyde and then stained with 0.2% crystal violet solution in 2% ethanol. The cells were counted under phase contrast inverted microscope.

### Migration Assay

Post 48 hours of transfection the cells (.1 million) were washed with PBS and resuspended in DMEM supplemented with 0.5% of FBS. Cells were then inoculated to the upper chamber of the transwell plate while in lower chamber DMEM supplemented with 0.5% FBS and 40μg/ml collagen I (Sigma-Aldrich, Missouri, USA) was added. After incubation for 24 hours in CO□incubator, the migrated cells were fixed with 5% glutaraldehyde for 20 mins and then stained with 0.2% crystal violet solution in 2% ethanol for 20 mins. The cells were counted under phase contrast inverted microscope.

### Real Time-qPCR

Both transfected and their control cells were harvested after 48 hours of transfection followed by total mRNA isolated using TRIZIN reagent (GCC Biotech, India). One step RT-qPCR kit (Invitrogen, Thermo Fisher Scientific, USA) was used according to manufacturer’s instructions to check the expression of Cdt2 at mRNA level by using the respective primers (forward and reverse^7^) mentioned in the Table 1.

**Table 1:**
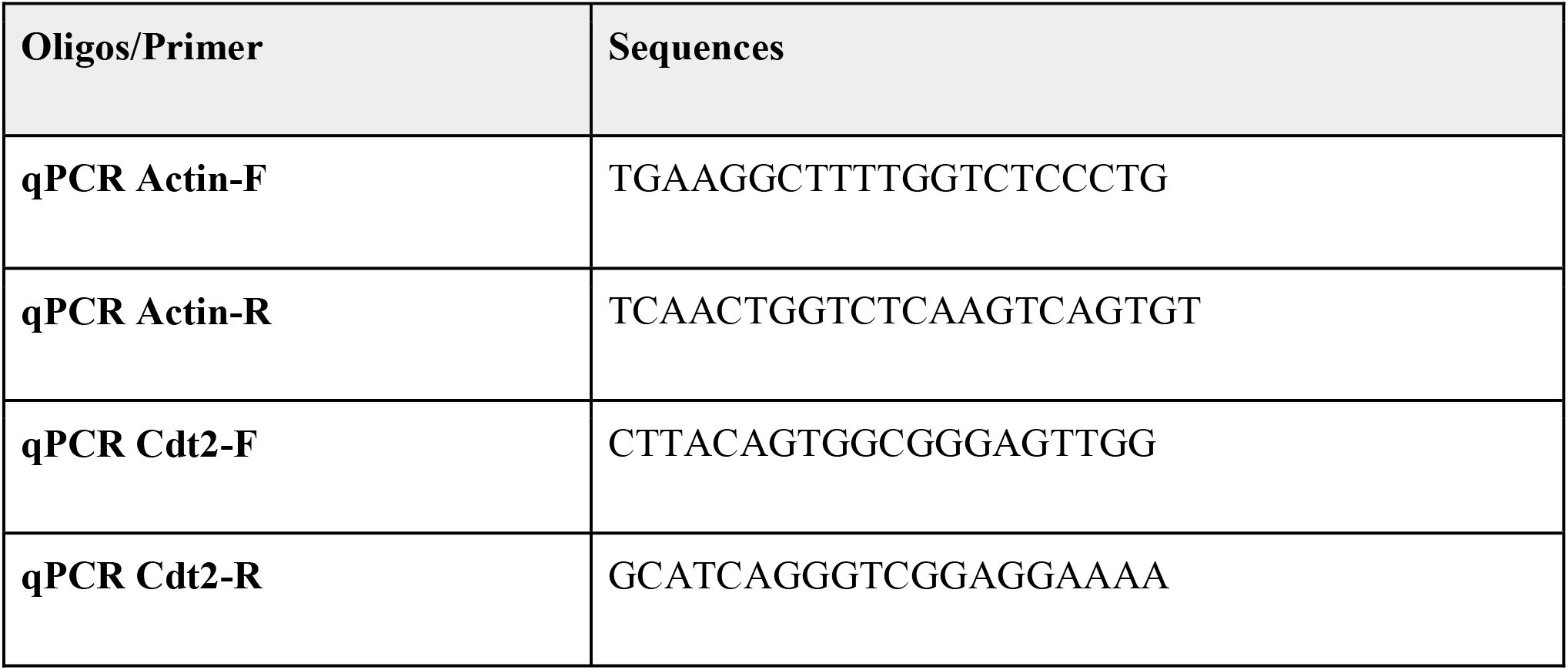
Sequences of primers

### Flow Cytometry

After transfection the cells were trypsinized and washed with PBS. The cells were then fixed in 70% ethanol at -20□overnight. Excess ethanol was washed with PBS followed by resuspension of the cells in 1 ml of staining solution (100 μg/ml propidium iodine, 50 mg/ml RNase-A in PBS and 0.1% Triton X-100)^29^. The stained cells were then analyzed using the flow cytometer (Beckman Coulter) the results were then compared and analyzed using the Cytoflex software provided with the flow cytometer system.

### Statistical Analysis

All the western experiments were performed in biological triplicates. The Flow cytometry, growth curve, migration and invasion assays were performed in experimental triplicates. All the data are presented as either mean ± SD or mean ± SE and an unpaired student’s t-test was performed to calculate the significant value. Image J software was used to quantify the intensity of protein bands wherever applicable.

## Results

### miR-17∼92 suppresses Cdt2 level in cervical cancer cells

Previous studies have shown that in cervical cancer cells level of Cdt2 is highly upregulated^7,8^. In an early screen we have found that miR-17∼92, a cluster of 6 mature micro RNAs (miR-17, miR-18a, miR-19a, miR-20a, miR-19b-1 and miR-92a-1) suppresses Cdt2 protein levels. In order to check the complementarity between miR-17∼92 cluster microRNA and Cdt2/DTL transcript we performed a microRNA databases search and we had found that according to miRdb, Micro T (Diana tool), miRwalk and RNA 22, multiple microRNAs (miR 20a-5p, miR17-5p, miR-17-3p, miR-18a, miR-92a) from the miR-17∼92 cluster interact with Cdt2/DTL transcript at 3’UTR (Figure 2A). To confirm the effect of miR-17∼92 cluster on Cdt2/DTL transcript in cervical cancer cells we transfected the miR-17∼92 in HPV positive cervical cancer cell SiHa, HPV negative cervical cancer cell, C33A and non-cancerous cell line HEK293T and observed that there is a significant decrease in transcript level of both HPV positive and negative cervical cancer cell lines but there is not much effect on Cdt2 transcript level in non-cancerous cell line HEK293T (Figure 2B). This confirms that miR17-92 directly binds to 3’ UTR and suppresses Cdt2 transcription. To check that if this effect is getting translated at protein level, we performed western blot for HPV positive and negative cervical cancer cells and discovered that there is a significant amount of decreases in the levels of Cdt2 protein in cancerous cells but no significant changes have been observed in non-cancerous cells (Figure 2C-2D).

**Figure 2:**
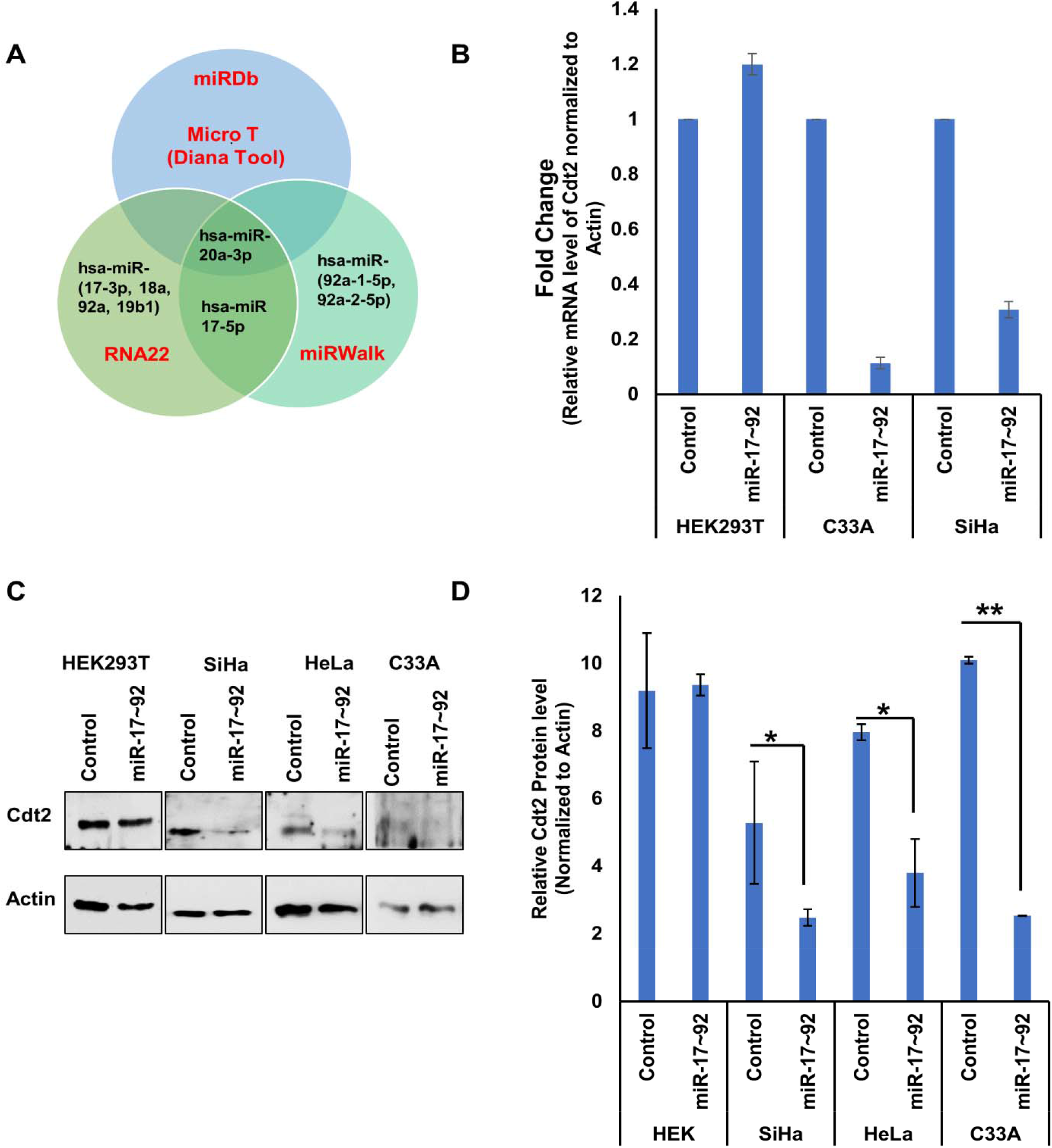
miR17-92 over expression decreases Cdt2 expression in cervical cancer cells. **A**. Different miRNAs from miR 17-92 cluster targets Cdt2/DTL according to different miRNA databases. **B**. RT-qPCR analysis of mRNA level of Cdt2 48h after transfection of miR-17∼92 in HEK293T, C33A and SiHa **C**. Western blot for analysis of Cdt2 protein expression level 48 hours after transfection of miR-17∼92 in HEK293T, SiHa, HeLa and C33A cell line. **D**. Quantification of Cdt2 protein level 48h post miR 17-92 transfection in HEK, SiHa, HeLa and C33A compared to vector control. Quantification experiments were done in triplicates. Error bar depicts standard error (S.E.). *p value <0.05

### miR-17∼92 blocks the cell cycle in S phase in cervical cancer cells

We have shown that miR-17∼92 suppresses Cdt2 which is a major S phase regulator, at transcript level in cervical cancer cells. In order to investigate the effect of miR-17∼92 on cell cycle progression, we ectopically expressed miR-17∼92 in SiHa cells and performed flow cytometry. We observed that there is an extensive cell cycle arrest at S phase i.e., there is an increase of population of cells from 12.4% to 27.4% in control versus miR-17∼92 treated cells, which is more than 2-fold increase. Also, miR-17∼92 moderately arrests cell cycle in G2/M phase and increases the cell population from 18.9% to 21.8% (Figure 3A). Additionally, we also observed accumulation of sub G0 population which represents the apoptotic cells.

**Figure 3:**
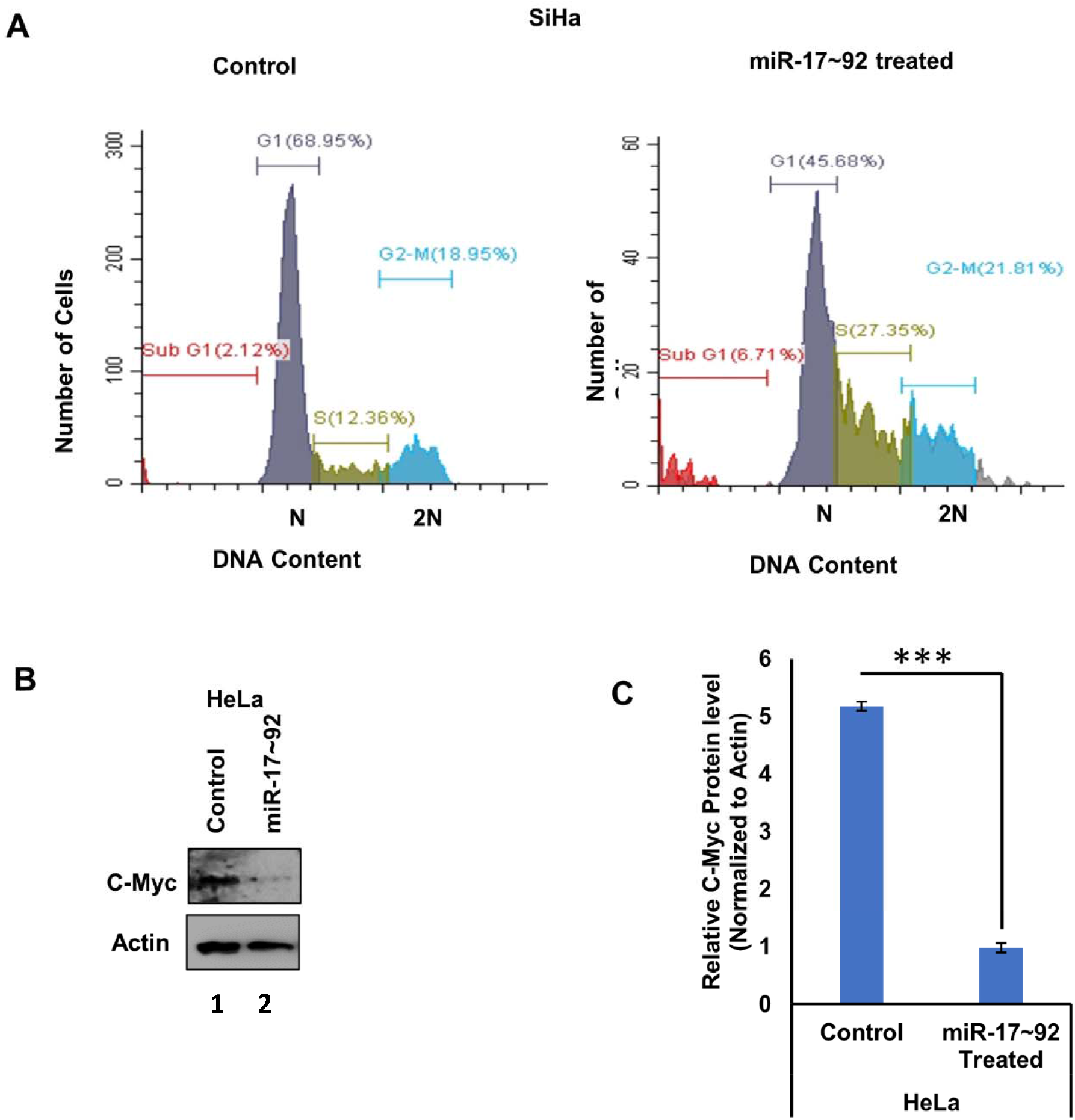
miR-17∼92 ectopic expression causes S phase arrest in cervical cancer cells. **A**. Flowcytometric analysis of SiHa cells post 4R hours of transfection of miR-17∼92. **B**. Western blot for analysis of C-Myc protein expression level 48 hours after transfection of miR-17∼92 in Hela Cells. **C**. Quantification of C-Myc protein level 48h post miR 17-92 transfection HcLa Cells compared to vector control. Quantification experiments were done in triplicates. Error bar depicts standard error (S.E.). ***p value <0.0001

Since C-Myc and miR-17∼92 cluster works in an autoregulatory loop, therefore next we wanted to confirm the effect of miR-17∼92 on C-Myc. As shown in previous literatures, we too confirmed that ectopic expression of miR-17∼92 cluster indeed significantly suppresses the level of C-Myc in cervical cancer cells (Figure 3B and 3C).

### miR-17∼92 suppresses the proliferation of cervical cancer cells

Since Cdt2 is one of the major player of cell proliferation in cervical cancer^15,30^ and we have already shown that miR-17∼92 suppresses Cdt2 level in cervical cancers cells, therefore next we wanted to check the effect of miR-17∼92 overexpression on growth and proliferation in cervical cancer cells. We observed that miR-17∼92 expression significantly inhibits the proliferation in HPV positive cervical cancer cells, HeLa and SiHa (Figure 4C-4F). Although there is suppression in growth in HPV negative cervical cancer cells C33A (Figure 4G and 4H) but there is no such effect on noncancerous HEK293T cells where the growth has decreased at 3^rd^ day and has been compensated by 4^th^ day (Figure 4A and 4B). Also, the phase contrast microscopy has shown that upon miR-17∼92 transfection the cervical cancer cells have incurred altered morphology and acquired a more rounded structure in comparison to wild type cell lines (Figure 3B, 3D, 3F and 3E) while no such change in the morphology of HEK293T cells has been observed upon transfection Figure 3B).

**Figure 4:**
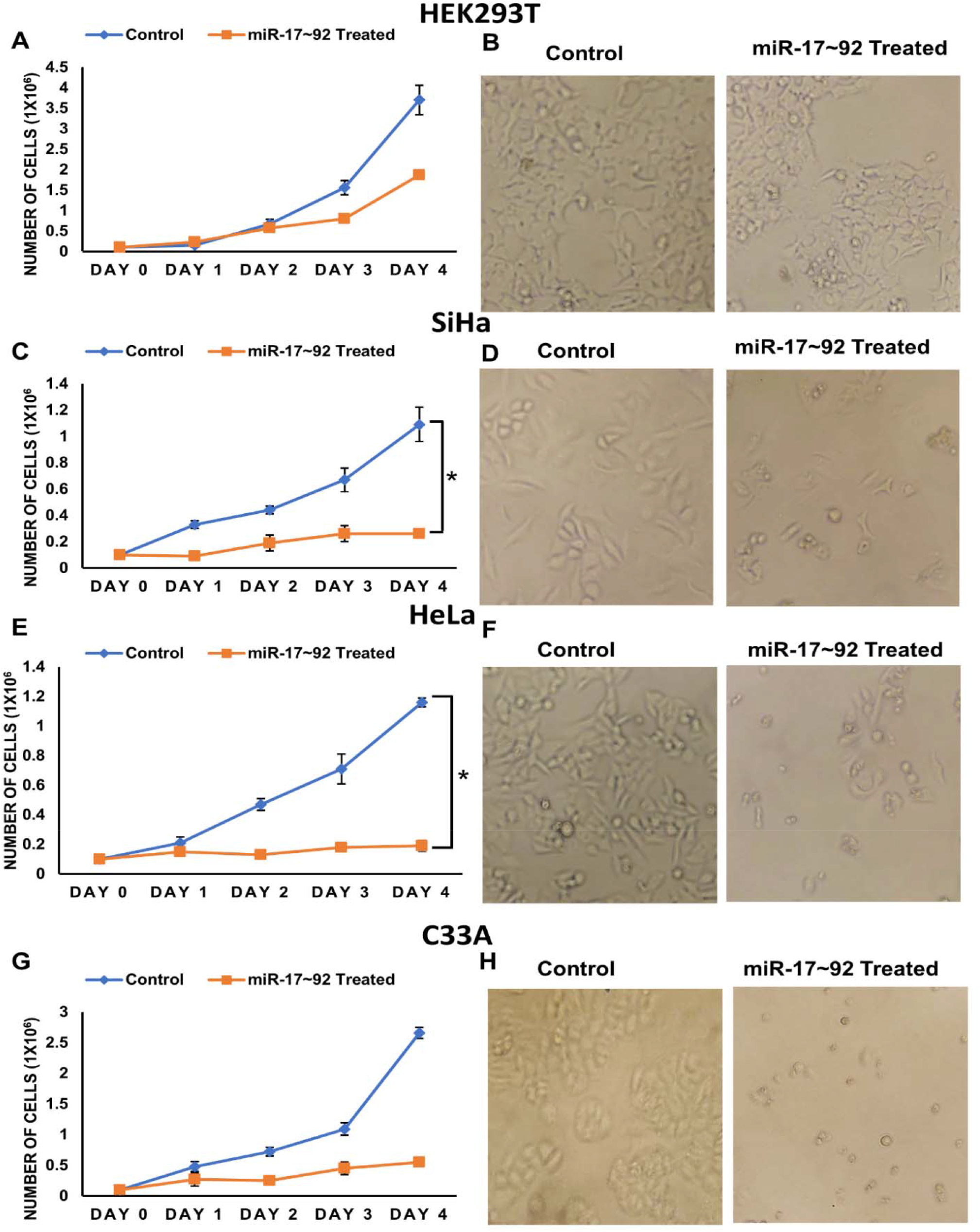
**miR17-92 suppresses proliferation of cervical cancer cells: miR17-92 ectopically expressed and the proliferation was observed from day of transfection till day 4 A. Growth curve of HEK293T cells with its respective control B. Control and treated HEK293T cells after 48h of transfection under phase contrast microscopy. C. Growth curve of SiHa along with its control D. SiHa cells (control and treated) under phase contrast microscopy after 48h of transfection. E. Growth curve of HeLa and it’s control F. Phase contrast microscopy of treated and control HeLa cells 48h post transfection. G. Growth curve of C33A and its control after transfection F. C33A control and treated cells under phase contrast microscopy after 8h of transfection. The experiment was done in triplicate. Error bar represents S.D. *p value-<0.05, **p value<0.001**

### Upregulation of miR-17∼92 suppresses both cell migration and invasion of cervical cancer cell lines

After establishing that miR-17∼92 inhibits proliferation in cervical cancer cells next, we checked the effect of miR-17∼92 on the cell migration and invasion ability of cervical cancer cells. The transwell invasion and migration assays has shown that miR-17∼92 reduces the invasion capability of HeLa cells by 34.2% and of SiHa cells by 40.5%.in comparison to the control (Figure 5A-5B). We also observed a suppression in migration ability of HeLa cells by ∼0.7 fold and that in SiHa cells by ∼0.9 fold (Figure 5C and 5D). Hence, invasion and migration ability of cervical cancer cells are significantly suppressed in cervical cancer cells upon miR17∼92 overexpression.

**Figure 5:**
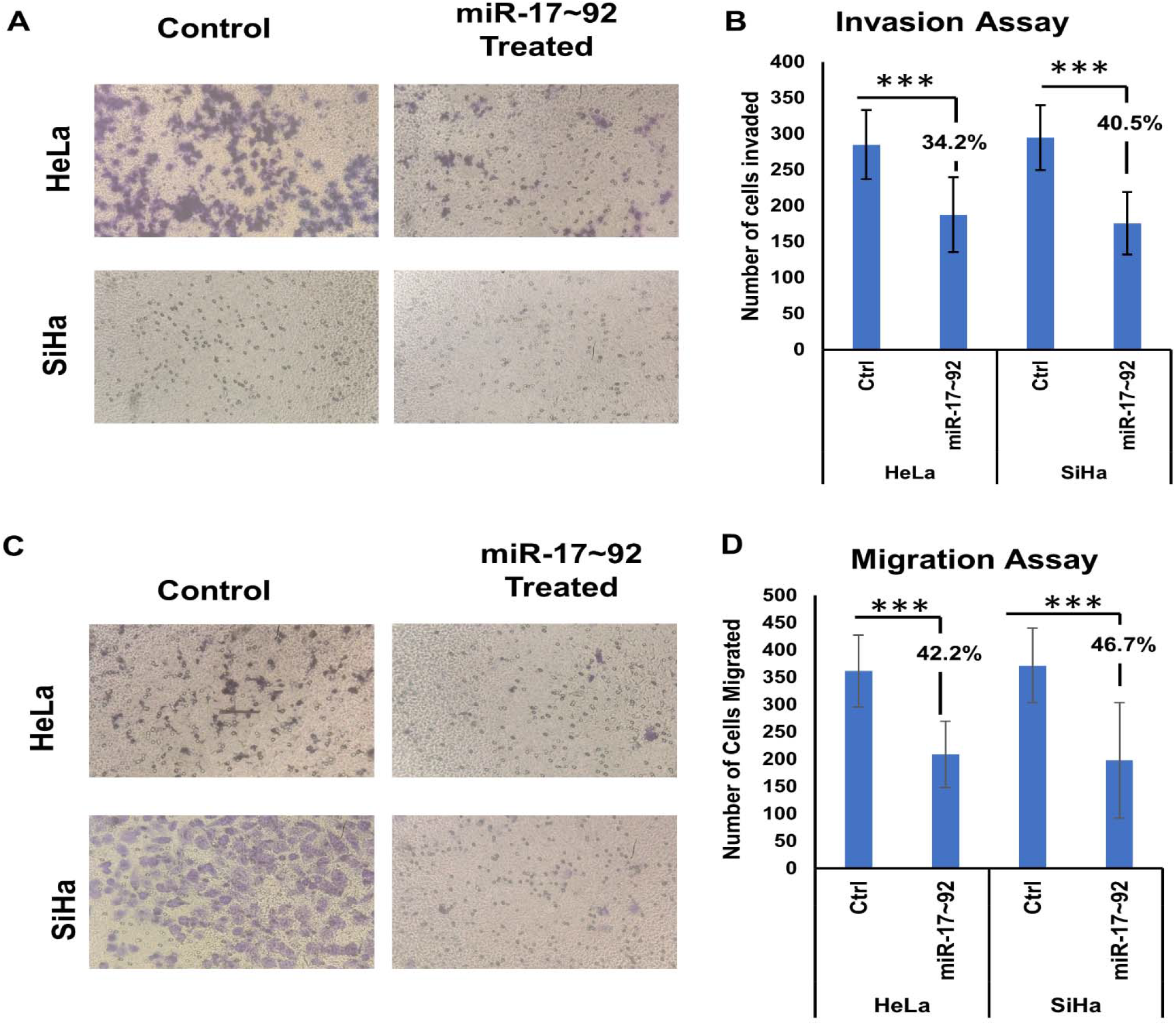
miR17-92 ectopic expression effect on transwell migration and invasion of cervical cancer cells **A**. Phase contrast microscopy images of control and treated SiHa and HeLa cells representing the change in invasion ability of treated cells against their respective controls **B**. Graphical representation of invasion assay of HeLa and SiHa cells along with their control **C**. Images representing the migration of treated HeLa and SiHa along with their controls. **D**. Graphical representation of migration assay of HeLa and SiHa cells along with their control. 8 different areas were selected randomly and number of cells invaded and migrated cells were counted. Error bar depicts standard deviation (S.D.). *p value <0.0001

## Discussion and Conclusion

Cervical cancer is fourth most aggressive and fatal cancer among women worldwide associated with the mutation and deletion in miRNA genes and their aberrant expression^3,31^. One such miRNA is miR-17∼92 cluster (encoded by 13q31.3) which consist of 6 mature miRNAs: miR-17, miR-18a, miR-19a, miR-20a, miR-19b-1 and miR-92a-1, displaying both tumor suppressor and oncogenic functions and works in an autoregulatory manner in order to maintain the growth and proliferation of the cells^19–21^. In cervical cancer cases specifically in HPV infected one, E6 protein apart from affecting other pathways also modulate the expression of miRNA-92 from the cluster and increase the proliferation of the infected cells^32^. However, nothing much is known about mechanism behind regulation of cell cycle factors by miR-17-92 in cervical cancers.

In this study, we show that upon over expression of miR-17∼92 it suppresses the proliferation of cervical cancer cells. To confirm the tumor suppressor role of miR-17∼92 we performed the miRNA database search using bioinformatic tools to identify the potential targets of miR-17∼92 cluster and found out that several miRNAs from cluster indeed suppresses an essential cell cycle factor Cdt2/DTL which is involved in G1 to S phase transition^10,33,34^ at the 3’UTR (Figure 2A). This in-silico information was validated by RT-qPCR where we have shown that miR-17∼92 suppresses the mRNA level of Cdt2 both in HPV negative and positive cervical cancer cells but there was no suppression in non-cancerous cells (Figure 2B). On the basis of complementarity of miRNA seed sequences to the 3’UTR of the target mRNA, miRNAs have the ability to either degrade the mRNA or inhibit the translation of protein^16^. This indicates that miR-17-92 might be a natural suppressor of Cdt2 but since Cdt2 levels are quite higher in CC cell lines, over expression of miR-17-92 regulates and bring back Cdt2 levels to normalcy which suppresses proliferation of these cancerous cells. This suppressive role of miR-17∼92 in the expression of Cdt2 was supported by the downregulation of Cdt2 protein only in cervical cancer cells (Figure 2C Lane 3-8 and 2D) but there was no significant change in non-cancerous cells (Figure 2C Lane 1-2 and 2D). This shows that suppressive nature of miR-17∼92 to Cdt2 is specific to cervical cancer cells only where Cdt2 has been reported to be upregulated.

Cdt2 is an adapter protein of CRL4 E3 ubiquitin ligase system which leads to the ubiquitin mediated degradation of cell cycle pro-apoptotic and onco-suppressor factors p21 and Set8 for the progression of cell from G1 to S phase and S to M phase^10,11,35^ which is in agreement with the flow cytometer result which shows that ectopic expression of miR-17∼92 in cervical cancer cell (SiHa) causes extensive S phase and moderate G2/M phase arrest.

It has been already reported that in cervical cancers Cdt2 is upregulated and is responsible for the proliferation and tumorigenesis^7^, therefore suppression of Cdt2 by miR-17∼92 shows significant reduction in growth and proliferation of HPV positive cervical cancer cells (Figure 4C-4F). Also, in HPV negative cells, C33A there is suppression in growth and proliferation although the suppression is not significant enough (Figure 4G-4H) whereas in HEK29T cells where Cdt2 level is not suppressed by miR-17∼92 expression the growth and proliferation remained unaffected. The other reason for the inhibition of growth is suppression of C-Myc level upon miR-17∼92 expression (Figure 3B and 3C). Because of the auto-regulatory nature of the miR-17∼92 via C-Myc regulation where miR-17∼92 suppresses the overly expressed C-Myc and suppresses the oncogenic signaling of C-Myc^19,20,23^. Additionally, miR-17∼92 overexpression inhibits the cell migration and invasion ability of cervical cancer cells (Figure 5A-5D) probably by inhibiting the proliferation of cells.

Cdt2 is an essential mammalian protein which couples with CRL4 ligase system as DECAF and form the CRL4Cdt2 complex which acts as the ubiquitin mediated degradation system for other cell cycle factor like p21 and Set8. In cervical cancer cells, Cdt2 level becomes upregulated and leads to the hyper-proliferation and re-replication. Our study for the first time shows that mir-17∼92 act as a tumor suppressor in cervical cancer cells by targeting Cdt2 and suppresses its mRNA and protein level which leads to the inhibition of proliferation, invasion and migration ability of cervical cancer cells by causing S phase cell cycle arrest (Figure 6) while there is no affect on the non-cancerous cells. Our study opens up a way to explore the possibility of using miR-17∼92 as cervical cancer treatment which could be much cheaper and specific therapeutic option for cervical cancer.

**Figure 6:**
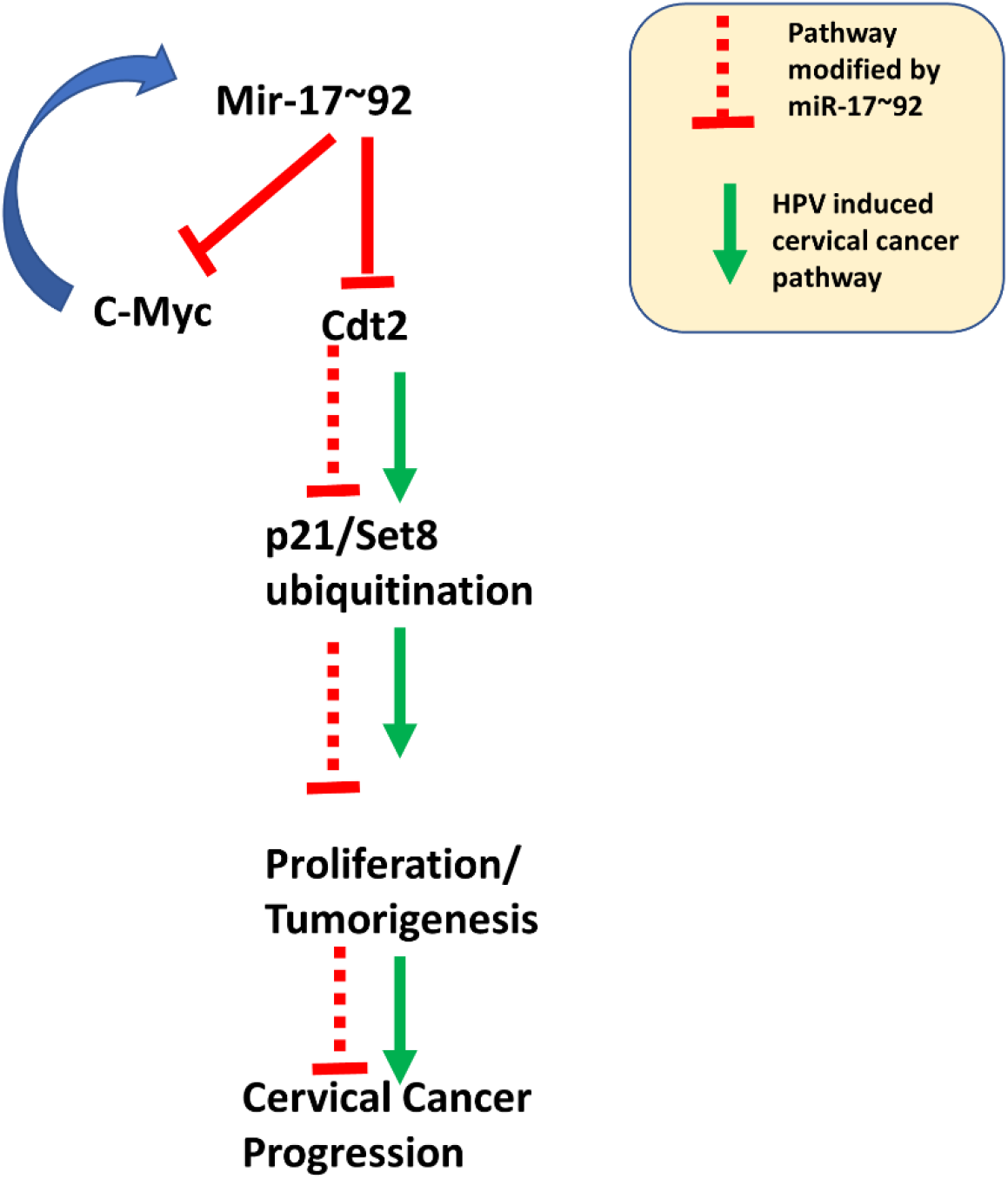
miR-17∼92 expression suppresses Cdt2 at mRNA level, leading to stabilization of p21 and Set8, ultimately decreases proliferation and tumorigenesis.

## Declarations

### Ethics approval and consent for participation

Not Applicable

### Consent for publication

Not Applicable

### Availibilty of data and materials

Not Applicable

### Competing interest

The authors declare no competing interest.

### Funding

The research was funded by Department of Biotechnology (DBT), Govt. of India, RLS grant (BT/RLF/Re-entry/43/2016) to Samarendra K Singh. The University Grants Commission (UGC), Govt. of India also supported this research by providing scholarship to Garima Singh, the Council of Scientific and Industrial Research (CSIR) by providing scholarship to Sonika Kumari Sharma and Department of Biotechnology (DBT) by providing scholarship to Astha (M.Sc. Stipend).

## Authors’ contributions

GS performed all the major experiments and did data analysis. She was also involved in writing, reviewing and editing of the manuscript. SKS and Aastha helped GS in the experiments and data analysis and was involved in writing and editing of the manuscript. SKS was involved in the conceptualization, designing, supervision, writing, critical review and editing of the manuscript.

## Acknowledgement

The authors are thankful to the Director Prof. A.K. Tripathi, School of Biotechnology, Institute of Science, Banaras Hindu University for providing space and facilities to conduct the research. We thank the Central Discovery Center (CDC) for facilitating the chemidoc and flow cytometry facility. We are thankful to Department of Biotechnology, Govt. of India for funding Samarendra K Singh (SKS) and Aastha, University Grants Commission (UGC), Govt. of India for funding Garima Singh and the Council of Scientific and Industrial Research (CSIR), New Delhi for funding Sonika Kumari Sharma.

## Notes

### Competing Interest Statement

The authors have declared no competing interest.

